# Glycowork: A Python package for glycan data science and machine learning

**DOI:** 10.1101/2021.04.22.440981

**Authors:** Luc Thomès, Rebekka Burkholz, Daniel Bojar

## Abstract

As a biological sequence, glycans occur in every domain of life and comprise monosaccharides that are chained together to form oligo- or polysaccharides. While glycans are crucial for most biological processes, existing analysis modalities make it difficult for researchers with limited computational background to include information from these diverse and nonlinear sequences into standard workflows. Here, we present glycowork, an open-source Python package that was designed for the processing and analysis of glycan data by end users, with a strong focus on glycan-related data science and machine learning. Glycowork includes numerous functions to, for instance, automatically annotate glycan motifs and analyze their distributions via heatmaps and statistical enrichment. We also provide visualization methods, routines to interact with stored databases, trained machine learning models, and learned glycan representations. We envision that glycowork can extract further insights from any glycan dataset and demonstrate this with several workflows that analyze glycan motifs in various biological contexts. Glycowork can be freely accessed at https://github.com/BojarLab/glycowork/.

## Introduction

Discovering trends and patterns in biological data requires i) large datasets and ii) dedicated algorithms from data science, bioinformatics, or machine learning. The combination of these two factors has led to great advances in systems biology (Chuang, Hofree and Ideker 2010; Zou and Laubichler 2018). Usually, there is limited overlap between groups engaging in data collection and those developing algorithms. In mature systems biology fields, this is abrogated by user-friendly software that facilitates mostly experimental end users to analyze their datasets for routine applications, such as with the Bioconductor (Huber *et al.* 2015) or Biopython (Cock *et al.* 2009) platforms.

Glycobiology – the analysis of glycans and the roles of these complex carbohydrates in biological contexts (Varki 2017) – has recently seen a surge in both data gathering and algorithmic development. Moderately large datasets from sources such as glycomics (Cummings and Pierce 2014), glycan arrays (Oyelaran and Gildersleeve 2009), or lectin arrays (Ribeiro and Mahal 2013) can by now be gathered on a more or less routine basis, depending on the application. Many algorithms for the analysis of these glycan-related data, such as subtree mining for the analysis of glycan array data (Coff *et al.* 2020; Haab and Klamer 2020) or glycan-focused machine learning (Bojar *et al.* 2021; Burkholz, Quackenbush and Bojar 2021), have been recently developed. Yet while both factors that are necessary for effective analysis – data and algorithms – are present in glycobiology, most efforts of algorithm development in the field are inaccessible to the typical end user, who may not be well-versed in computational workflows. While webtools and graphical user interfaces for some applications have been developed (Grant *et al.* 2016; Huang *et al.* 2021) and are considerably more accessible, these approaches often lack the flexibility and throughput that is required for many analyses. Thus, with the exception of platforms such as glypy (Klein and Zaia 2019), geared more toward the analysis of glycan-focused mass spectrometry, glycobioinformatics methods cannot be used with the same ease and accessibility that bioinformatics procedures exhibit in other systems biology disciplines.

For this purpose, we have developed glycowork, a computational framework that is designed to be accessible and usable by end users with minimal computational background. While the background functions that work with glycans as graphs are easily accessible by experts, we provide numerous high-level wrapper functions that only require the input of glycans in a human-readable format such as the IUPAC-condensed format and that allow end users to engage in motif analyses, plotting, sequence context analyses, and more. The code for glycowork is fully open-source (https://github.com/BojarLab/glycowork/) and we have prepared an extensive documentation with various example workflows for users to get started with this package (https://bojarlab.github.io/glycowork/). We envision glycowork to advance the spread and capabilities of glycobioinformatics, distilling insights from the increasing number of glycan datasets that are being gathered.

## Glycowork – Principles and applications

Glycowork is written in the Python programming language (version 3.6+) and uses pandas dataframes, lists of glycans, or single glycans as inputs for most of its functions. We have structured glycowork into four modules: data loading and handling (glycan_data), sequence alignment (alignment), sequence processing and motif analysis (motif), and glycan-focused machine learning (ml). All modules have submodules specialized for certain tasks (Fig. 1A).

**Figure 1.**
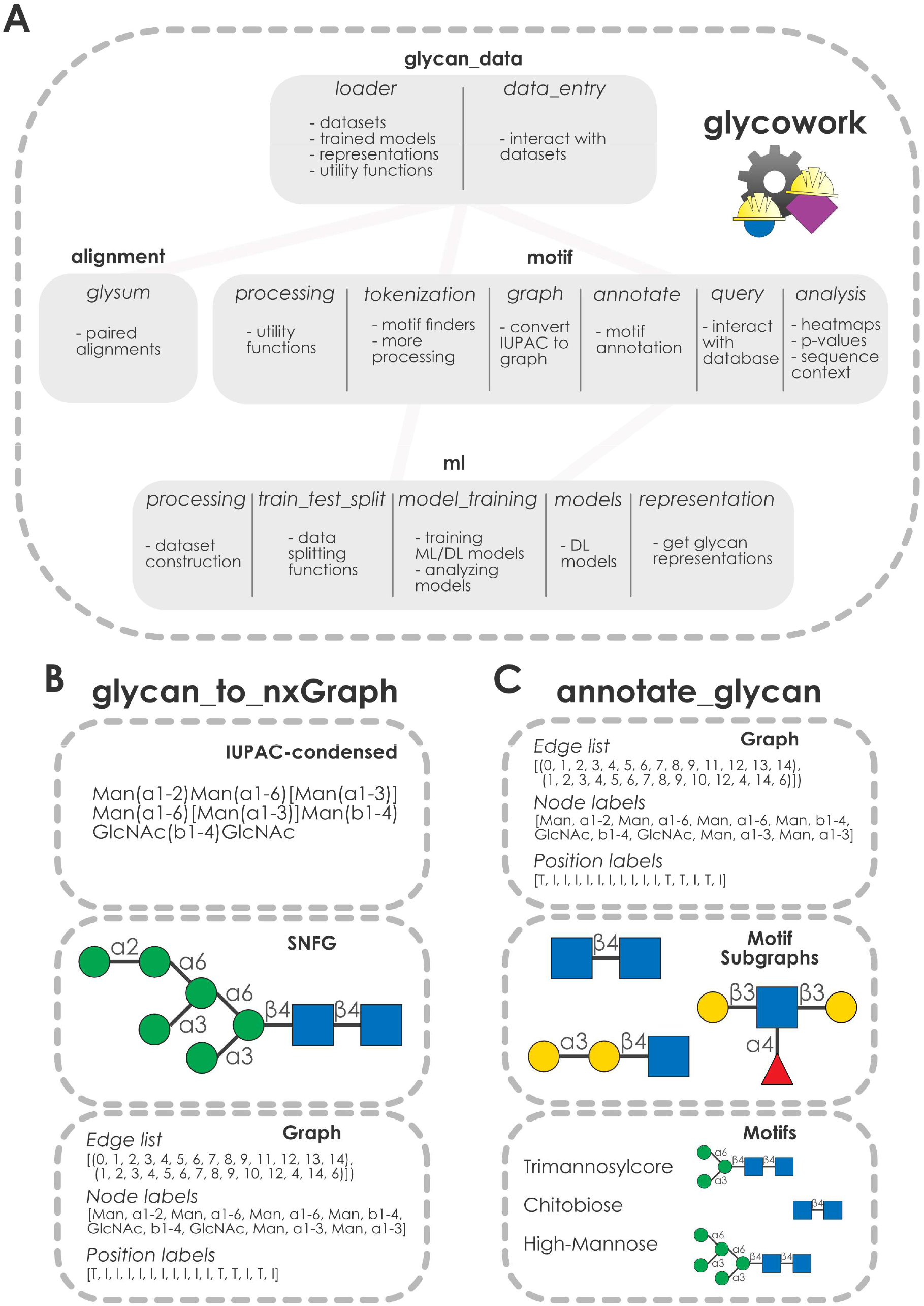
Structure of the glycowork package. **A)** Modular structure of glycowork. Modules are depicted as boxes with bold titles. Submodules within these modules are labelled with italics and example functionality is described for each submodule. Dependencies between modules are indicated by connecting lines. **B)** Workflow of the glycan_to_nxGraph function from the glycowork.motif.graph submodule. An example glycan in IUPAC-condensed notation is converted into a graph, capturing the graph-like nature of glycans depicted via the symbol nomenclature for glycans (SNFG). The resulting edge, node, and position lists are shown, with “T” indicating a terminal position and “I” indicating an internal position. **C)** Workflow of the annotate_glycan function from the glycowork.motif.annotate submodule. Glycan graph objects and graphs for various known motifs are used to identify occurring motifs (shown in the bottom box) via subgraph isomorphism tests.

Functions in glycowork routinely use glycans in the IUPAC-condensed nomenclature as input and convert these to graph objects (Burkholz, Quackenbush and Bojar 2021) that can be further processed and analyzed (Fig. 1B). Crucially, this step removes the ambiguity of the IUPAC nomenclature, as graph isomorphism can be explicitly tested. We decided against using other glycan nomenclatures, such as GlycoCT (Herget *et al.* 2008) or WURCS (Tanaka *et al.* 2014), in our workflows, as we argue that the combination of a human-readable nomenclature (IUPAC-condensed) with a machine-readable nomenclature (glycan graphs) is necessary and sufficient for all relevant tasks in glycobioinformatics. Further, working with glycans as general graphs allowed us to readily leverage advances in graph theory that have accrued over decades of research, such as present in the widely used NetworkX package (Hagberg, Schult and Swart 2008) that can be applied without modifications to our glycan graphs.

Glycan graphs consist of the connectivities present in a glycan structure (edge list), the contained monosaccharides / linkages (node labels), and the information whether these are internal or terminal in the sequence (position labels; Fig. 1B). While users can also input precomputed glycan graphs into most glycowork functions, the package is designed in a way that users with limited bioinformatics experience can exclusively work with glycans in IUPAC-condensed nomenclature, while all graph operations proceed smoothly in the background. We are confident that this will allow for a wide accessibility and adoption of our open-source package.

Many functions in glycowork explicitly leverage the power of graph theory, for instance by automatically and unambiguously annotating glycan motifs using the annotate_glycan function that detects motifs via subgraph isomorphism tests (Fig. 1C). This is done via positional matching, using the position labels, to ensure that motifs such as the *O*-glycan core motifs are only recognized if they occur at the reducing end. Glycowork comes equipped with a list of 150 named motifs from the academic literature and, for example, can also exhaustively analyze salient disaccharide motifs, as further described below.

Glycowork is equipped with continuously updated datasets of glycans, such as glycan array data of influenza viruses or information about species-specific glycans. These could be used for developing new algorithms, benchmarking existing algorithms, or uncovering new insights into properties of glycans or glycan motifs. Additionally, stored learned representations of glycans, derived from a deep learning model (Burkholz, Quackenbush and Bojar 2021), are provided. Together, this for instance allows us to visualize clusters of similar glycans that can be labeled with taxonomic properties or other labels (Fig. 2A). Further, glycowork contains functions to automatically annotate glycan motifs and cluster groups (for instance taxonomic groups) according to the presence and abundance of glycan motifs via heatmaps (Fig. 2A).

**Figure 2.**
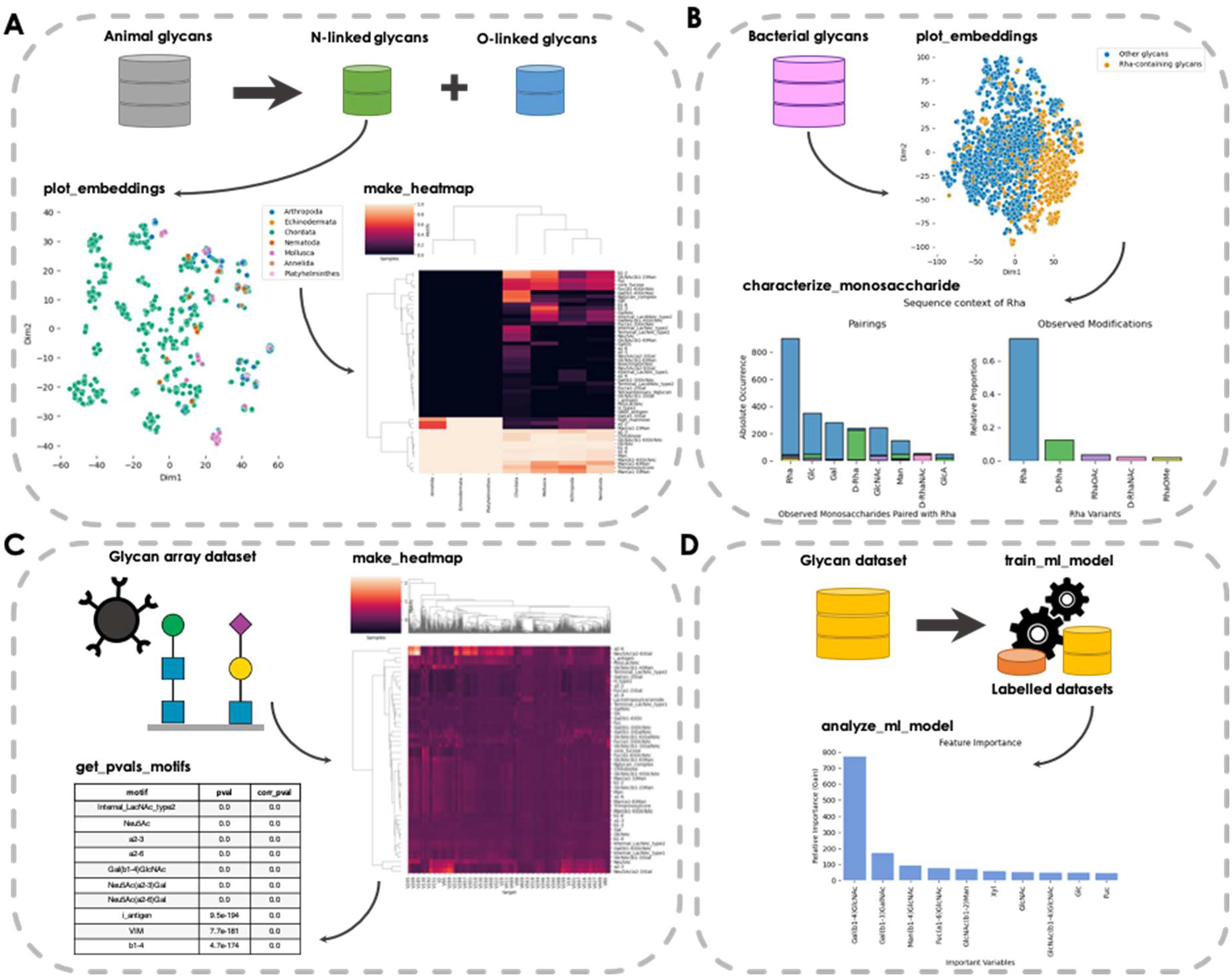
Example workflows from the glycowork package. **A)** Investigating N-linked glycans in animals. Dataset descriptions and names of glycowork functions are bolded. Data filtering and labelling processes are illustrated by large arrows and the order of the steps is indicated by small arrows. The plot_embeddings function of N-linked glycans displays each glycan as a colored dot, with colors corresponding to the taxonomic phyla. The heatmap is generated by the make_heatmap function and represents glycan motif distributions for each phylum. **B)** Analysis of rhamnose sequence context in bacteria. The diversity of bacterial glycans is depicted by the plot_embeddings graph. Dots are colored based on the presence of rhamnose (Rha) in each glycan. Rhamnose pairing preferences (left) and proportions of modifications (right) are visualized via the characterize_monosaccharide function. Pairing preferences of rhamnose variants are illustrated by stacked bar graphs. **C)** Glycan binding specificities of influenza viruses. Measured glycan binding of various influenza strains is represented as a heatmap for diverse motifs. The get_pvals_motifs function displays, for each motif, a p-value and a corrected p-value. Shown are the top ten motifs, with the full table available in Supplementary Table 1. **D)** Glycan classification using machine learning models. Glycans are first labelled with an “animal” or “non-animal” tag. Then, the train_ml_model function constructs a machine learning model to discriminate between these two classes. Finally, the analyze_ml_model function generates a bar graph to understand which criteria are most important for glycan classification. Full scale versions of the heatmaps shown in A and C can be found in Supplementary Fig. 1-2.

Besides these high-level analyses, users can engage in the analysis of the sequence context of a monosaccharide in a specific taxonomic group such as bacteria (Fig. 2B). Comparing the sequence context of different kingdoms or groups of interest might shed light onto evolutionary or functional differences in their glycans. Here, we show this type of analysis with the sequence neighborhood of the monosaccharide rhamnose (Rha) in bacteria. This yields the observation that Rha, D-Rha, and D-RhaNAc are all typically found in homogeneous sequence environments (i.e., connected to more Rha / D-Rha / D-RhaNAc, respectively).

Probing the glycan binding specificities of protein via glycan arrays is a common method to determine viral glycan-binding specificity (Smith and Cummings 2014). Glycowork can aid in the analysis of this type of data by generating motif-based heatmaps, in which motifs are colored by their associated Z-score – in this case illustrating the split between Neu5Ac(α2-3)-binding avian influenza viruses and Neu5Ac(α2-6)-preferring mammalian influenza viruses (Fig. 2C). This can be further extended by identifying statistically significant binding motifs from these data, a necessary step in determining viral binding motifs (Fig. 2C), which again points to the importance of sialic acid-containing motifs as reported previously (Viswanathan *et al.* 2010).

Another example of the functionalities found in glycowork is glycan-focused machine learning. By providing a list of glycans and corresponding labels, glycowork allows the training of machine learning models with a single line of code. As an example, we trained a model to predict whether a glycan stems from an animal or a different organism and then analyzed the model as to which motifs were most predictive for this classification (Fig. 2D). In this case, the presence of type-2 LacNAc (Gal(β1-4)GlcNAc) was most predictive for an animal glycan. Analogously, state-of-the-art deep learning models can be trained with only a few lines of codes using glycowork, for which we direct the user to the full documentation of glycowork.

## Conclusion

As the most diverse biological sequence, glycans are especially in need of dedicated analysis platforms. Until now, technical limitations have largely directed the focus of researchers on rather short and / or uniform glycans that are amenable to manual analysis, such as *N*-glycans or short *O*-glycans. Yet with the addition of more and more sequences (Malaker *et al.* 2021) – as well as more and more complex sequences – and the combination of glycan data with systems biology data (Kearney *et al.* 2021), manual analysis is becoming increasingly unrealistic. We envision that the ability of accessible analysis platforms such as glycowork will allow researchers to connect glycobiology knowledge from different areas to fuel discoveries of new trends and patterns as well as extend the scope of already known phenomena.

Glycowork is designed to be modular as well as extendable and we are planning to improve it in future work by expanding its functionalities. One direction will be to include implementations of more existing glycobioinformatics techniques, such as multiple glycan alignments using the Multiple Carbohydrate Alignment with Weights (MCAW) tool (Hosoda, Akune and Aoki-Kinoshita 2017). We will also update the stored datasets in glycowork as new glycans become available, to maximize the utility of functions for sequence context analysis, database queries, and others.

We encourage interested readers to find more details, implementations, and examples in the documentation of glycowork (https://bojarlab.github.io/glycowork/). We also would like to invite the community to suggest – or even implement – changes, improvements, or additions, to maximize the utility of glycowork for glycobioinformatics and allow researchers to include glycan data analysis into their routine workflows.

## Acknowledgments

The authors would like to thank Frédérique Lisacek and Benjamin P. Kellman for helpful discussions.

## Author Contributions

Conceptualization: D.B., Data Curation: L.T., R.B., D.B., Funding Acquisition: D.B., Investigation: L.T., D.B., Resources: D.B., Software: L.T., R.B., D.B., Supervision: D.B., Visualization: L.T., D.B., Writing – Original Draft Preparation: L.T., D.B., Writing – Review & Editing: L.T., R.B., D.B.

## Funding

This work was funded by a Branco Weiss Fellowship – Society in Science awarded to D.B., by the Knut and Alice Wallenberg Foundation, and the University of Gothenburg, Sweden.

## Declaration of Interests

The authors declare no competing interests.

## Data Availability Statement

All code and data used for this work can be freely accessed at https://github.com/BojarLab/glycowork/

## Notes

https://bojarlab.github.io/glycowork/

### Competing Interest Statement

The authors have declared no competing interest.

