## Supplemental Figures for "Glycowork: A Python package for glycan data science and machine learning"

**Supplementary Figures**

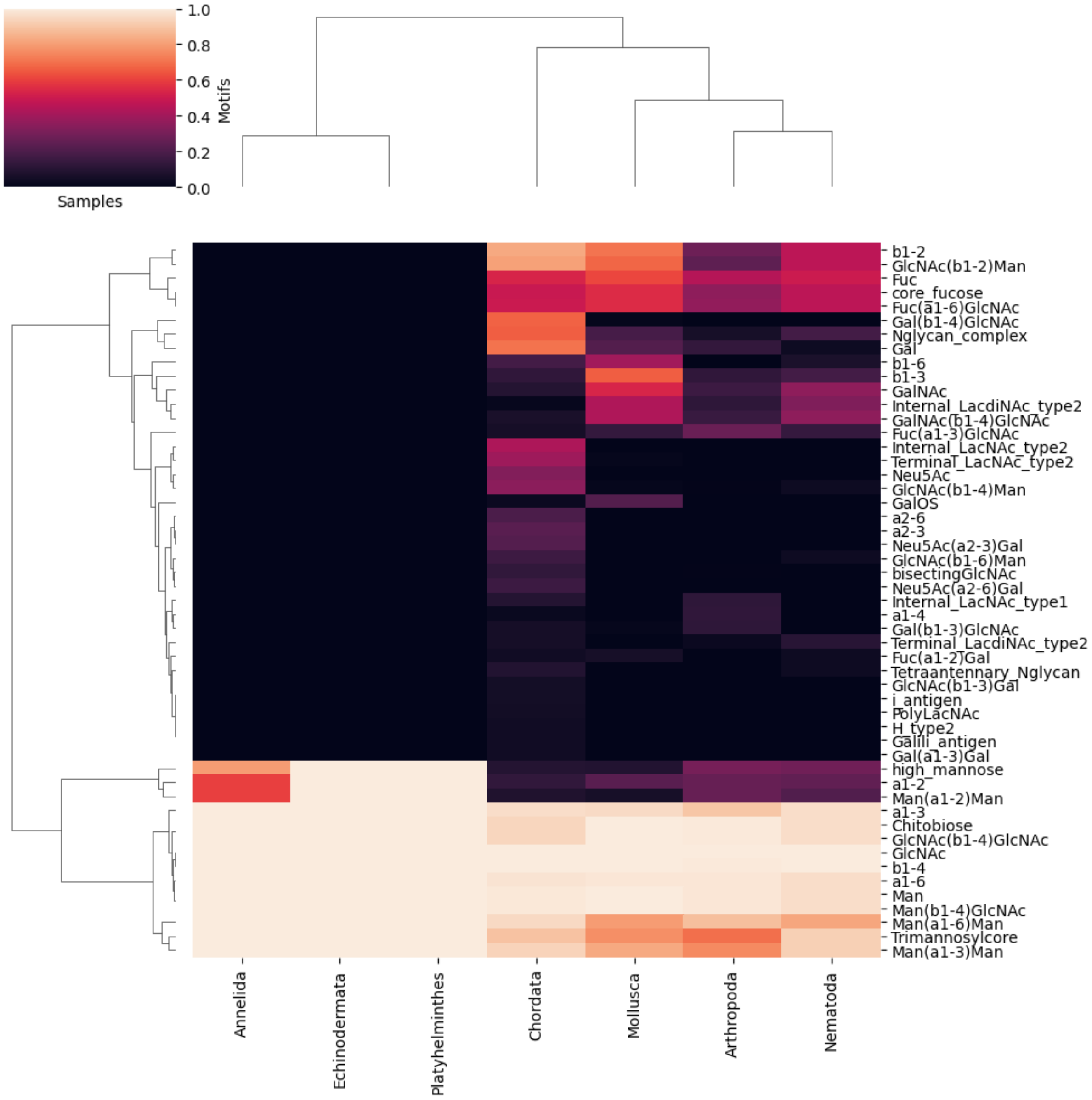


**Supplementary Figure 1. Heatmap of glycan motif distribution among animal phyla.** Related to Figure 2A. The heatmap is generated using the make_heatmap function and represents glycan motif distributions from animal N-linked glycans. Colors indicate the relative abundance of glycan motifs in each phylum. Motifs are labeled and correspond either to monosaccharides, linkages, disaccharides, or to larger structures such as polylactosamine repeats (PolyLacNAc).

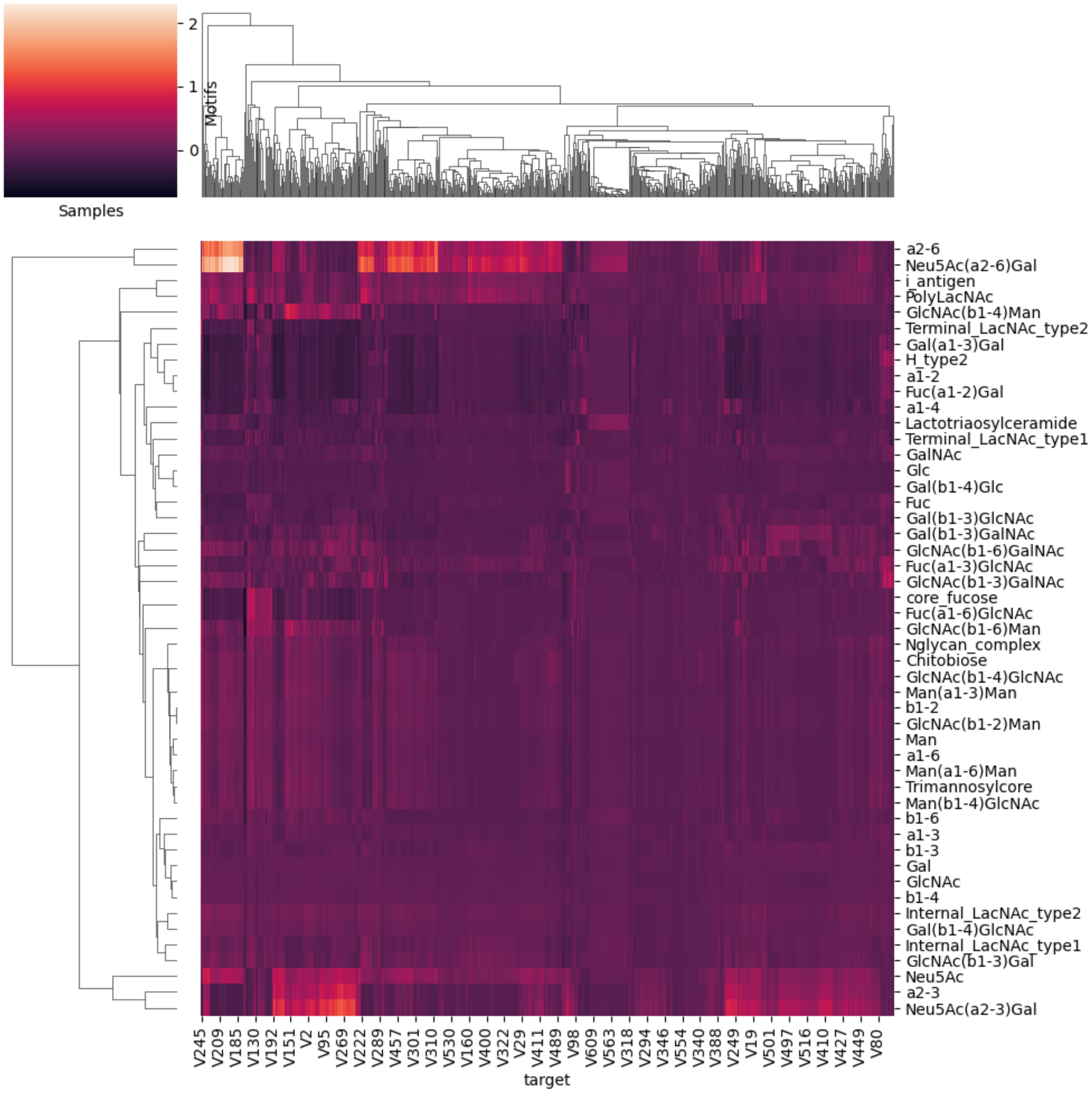


**Supplementary Figure 2. Motif-based heatmap of glycan binding specificities between influenza strains.** Related to Figure 2C. The heatmap is generated by the make_heatmap function and displays the measured glycan binding efficiency (Z-score) of various influenza strains for diverse motifs. The color gradient represents Z-scores associated with a motif. Motifs are labeled as monosaccharides, linkages, disaccharides, or larger structures. Influenza strains are represented by their identifiers.

### Supplementary Tables

**Supplementary Table 1. Statistical significance of glycan binding specificities of influenza viruses.** Related to Figure 2C. This table corresponds to the full table generated using the get_pvals_motifs function. The pval and corr_pval columns contain p-values and corrected p-values (corrected by the Holm-Šídák method) respectively, which represent the statistical significance for each binding motif. Columns are sorted ascendingly by p-values.

| motif | pval | corr_pval |
|---|---|---|
| **Internal_LacNAc_type2** | 0.0 | 0.0 |
| **Neu5Ac** | 0.0 | 0.0 |
| **a2-3** | 0.0 | 0.0 |
| **a2-6** | 0.0 | 0.0 |
| **Gal(b1-4)GlcNAc** | 0.0 | 0.0 |
| **Neu5Ac(a2-3)Gal** | 0.0 | 0.0 |
| **Neu5Ac(a2-6)Gal** | 0.0 | 0.0 |
| **i_antigen** | 9.53871705085845e-194 | 0.0 |
| **VIM** | 7.690771572918131e-181 | 0.0 |
| **b1-4** | 4.671886339472552e-174 | 0.0 |
| **PolyLacNAc** | 4.703869237162888e-155 | 0.0 |
| **Gal(b1-4)GlcNAcOS** | 1.0314895321484356e-108 | 0.0 |
| **GlcNAc(b1-3)Gal** | 2.761773896229227e-106 | 0.0 |
| **Gal** | 1.1530767361645484e-94 | 0.0 |
| **GlcNAc** | 1.1703728505419271e-86 | 0.0 |
| **GlcNAcOS** | 1.8129332905909683e-86 | 0.0 |
| **SialylLewisX** | 2.8909637444671393e-60 | 0.0 |
| **GlcNAc(b1-2)Man** | 1.1025544114971412e-59 | 0.0 |
| **Neu5Ac(a2-6)GalNAc** | 2.779528553149326e-59 | 0.0 |
| **b1-2** | 6.652734627105335e-56 | 0.0 |
| **Fuc_LN3** | 6.652681483864124e-46 | 0.0 |
| **Oglycan_core2** | 6.717992600099721e-46 | 0.0 |
| **Internal_LacNAc_type1** | 5.155571589264967e-43 | 0.0 |
| **bisectingGlcNAc** | 3.562003433491632e-37 | 0.0 |
| **Chitobiose** | 2.835833054849996e-36 | 0.0 |
| **GlcNAc(b1-4)GlcNAc** | 2.835833054849996e-36 | 0.0 |
| **LexLex** | 2.096418371667195e-34 | 0.0 |
| **Man(a1-3)Man** | 1.7473001542302298e-33 | 0.0 |
| **Gal(b1-3)GalNAcOS** | 3.923738240471641e-33 | 0.0 |
| **Neu5Ac(a2-6)GlcNAc** | 6.491293865539391e-33 | 0.0 |
| **Disialyl_T_antigen** | 5.778262296664894e-32 | 0.0 |
| **Neu5Ac(a2-3)GalOS** | 3.8889908326093376e-31 | 0.0 |

| | | |
|---|---|---|
| **Nglycan_complex2** | 1.328706611653617e-30 | 0.0 |
| **Man(b1-4)GlcNAc** | 2.7416848220399755e-28 | 0.0 |
| **Internal_LacdiNAc_type2** | 7.054007106134345e-28 | 0.0 |
| **Man(a1-3)GalNAc** | 7.876721632785831e-28 | 0.0 |
| **Man(a1-6)GalNAc** | 7.876721632785831e-28 | 0.0 |
| **GlcNAc(b1-4)Man** | 1.6059572572004042e-23 | 0.0 |
| **Trimannosylcore** | 1.661513107391722e-23 | 0.0 |
| **SDLex** | 7.71176740084554e-23 | 0.0 |
| **Nglycan_complex** | 1.6189628923989955e-22 | 0.0 |
| **Fuc(a1-3)GlcNAcOS** | 8.444679157958654e-22 | 0.0 |
| **GalNAcOS** | 2.5656441621709624e-20 | 0.0 |
| **VIM2** | 8.025104518698597e-20 | 0.0 |
| **GlcNAc(b1-6)GalNAc** | 9.802382312005959e-20 | 0.0 |
| **Gal(b1-3)Neu5Ac** | 7.752261303193785e-18 | 0.0 |
| **SialylLewisA** | 2.988068353502674e-16 | 7.127631818093505e-14 |
| **GalNAc(b1-4)GlcNAc** | 6.198036366270052e-15 | 1.3242740237728867e-12 |
| **Tetraantennary_Nglycan** | 1.9453714871974113e-12 | 4.1241055015461825e-10 |
| **Man(a1-6)Man** | 4.836313046316612e-12 | 1.0204705969130146e-09 |
| **GalOS(b1-4)GlcNAc** | 6.381181214224658e-11 | 1.3400487275383455e-08 |
| **Oglycan_core4** | 1.5171044957836459e-10 | 3.170747764347226e-08 |
| **3SGb5** | 4.748624064051575e-09 | 9.877133169133856e-07 |
| **Gal(b1-3)GlcNAcOS** | 6.8614375221294304e-09 | 1.420316564804125e-06 |
| **Fuc(a1-3)GlcNAc** | 9.90774947281499e-09 | 2.04099432632443e-06 |
| **GlcNAc(b1-6)Man** | 1.0499359020091819e-08 | 2.152366299967845e-06 |
| **Internal_LacdiNAc_type1** | 1.5451739230697985e-07 | 3.152105367087987e-05 |
| **O_linked_mannose** | 4.71811168309478e-06 | 0.0009573204072889085 |
| **Gal(b1-3)GalNAc** | 1.2858455438171447e-05 | 0.0025940543091963475 |
| **b1-3** | 1.4242938188232921e-05 | 0.002858756913576621 |
| **GalNAc(b1-4)GlcNAcOS** | 3.767002993218919e-05 | 0.0075058373441107 |
| **Neu5Ac(a2-3)GalNAc** | 0.00020169042885934986 | 0.03934548795688764 |
| **GD1a** | 0.0002554733662579807 | 0.04933181502348283 |
| **cisGM1** | 0.001935522473804221 | 0.31727791769295943 |
| **b1-6** | 0.004427371372422693 | 0.5809191575428146 |
| **Fuc(a1-3)Gal** | 0.00950655484692593 | 0.8447387292261651 |
| **Man** | 0.07213014831453363 | 0.9999995074045557 |
| **GlcNAcOS(b1-3)GalOS** | 0.10862369038043973 | 0.9999999997699671 |
| **LewisX** | 1.0 | 1.0 |
| **LewisY** | 1.0 | 1.0 |
| **LewisA** | 1.0 | 1.0 |
| **LewisB** | 1.0 | 1.0 |
| **H_type2** | 1.0 | 1.0 |

| | | |
|---|---|---|
| **H_type1** | 1.0 | 1.0 |
| **A_antigen** | 1.0 | 1.0 |
| **Galili_antigen** | 1.0 | 1.0 |
| **GloboH** | 1.0 | 1.0 |
| **Gb5** | 1.0 | 1.0 |
| **Gb4** | 1.0 | 1.0 |
| **Gb3** | 1.0 | 1.0 |
| **Forssman_antigen** | 1.0 | 1.0 |
| **iGb3** | 1.0 | 1.0 |
| **I_antigen** | 1.0 | 1.0 |
| **Terminal_LacNAc_type1** | 1.0 | 1.0 |
| **Terminal_LacNAc_type2** | 1.0 | 1.0 |
| **Terminal_LacdiNAc_type1** | 1.0 | 1.0 |
| **Terminal_LacdiNAc_type2** | 1.0 | 1.0 |
| **Ganglio_Series** | 1.0 | 1.0 |
| **Lacto_Series(LewisC)** | 1.0 | 1.0 |
| **NeoLacto_Series** | 1.0 | 1.0 |
| **LewisD** | 1.0 | 1.0 |
| **Lactotriaosylceramide** | 1.0 | 1.0 |
| **GM3** | 1.0 | 1.0 |
| **H_type3** | 1.0 | 1.0 |
| **GM2** | 1.0 | 1.0 |
| **GM1** | 1.0 | 1.0 |
| **GD3** | 1.0 | 1.0 |
| **GD2** | 1.0 | 1.0 |
| **GD1b** | 1.0 | 1.0 |
| **Nglycolyl_GM2** | 1.0 | 1.0 |
| **GT1b** | 1.0 | 1.0 |
| **GD1** | 1.0 | 1.0 |
| **GD1a_2** | 1.0 | 1.0 |
| **GT3** | 1.0 | 1.0 |
| **GT2** | 1.0 | 1.0 |
| **GT1c** | 1.0 | 1.0 |
| **2Fuc_GM1** | 1.0 | 1.0 |
| **GQ1b** | 1.0 | 1.0 |
| **O_mannose_Lex** | 1.0 | 1.0 |
| **core_fucose** | 1.0 | 1.0 |
| **Isoglobotetraosylceramide** | 1.0 | 1.0 |
| **polySia** | 1.0 | 1.0 |
| **high_mannose** | 1.0 | 1.0 |
| **Gala_series** | 1.0 | 1.0 |

| | | |
|---|---|---|
| **Oglycan_core3** | 1.0 | 1.0 |
| **Oglycan_core5** | 1.0 | 1.0 |
| **Oglycan_core6** | 1.0 | 1.0 |
| **Sialosylparagloboside** | 1.0 | 1.0 |
| **LDNF** | 1.0 | 1.0 |
| **4dGal** | 1.0 | 1.0 |
| **4dNeu5Ac** | 1.0 | 1.0 |
| **6dGal** | 1.0 | 1.0 |
| **7dNeu5Ac** | 1.0 | 1.0 |
| **8dNeu5Ac** | 1.0 | 1.0 |
| **9dNeu5Ac** | 1.0 | 1.0 |
| **Fuc** | 1.0 | 1.0 |
| **GalNAc** | 1.0 | 1.0 |
| **GalOP** | 1.0 | 1.0 |
| **GalOS** | 1.0 | 1.0 |
| **Glc** | 1.0 | 1.0 |
| **GlcA** | 1.0 | 1.0 |
| **GlcN** | 1.0 | 1.0 |
| **GlcNAcOP** | 1.0 | 1.0 |
| **GlcNAcOProp** | 1.0 | 1.0 |
| **GlcOS** | 1.0 | 1.0 |
| **GlcOSA** | 1.0 | 1.0 |
| **Kdn** | 1.0 | 1.0 |
| **MurNAc** | 1.0 | 1.0 |
| **Neu** | 1.0 | 1.0 |
| **Neu5Ac9Ac** | 1.0 | 1.0 |
| **Neu5AcOMe** | 1.0 | 1.0 |
| **Neu5Gc** | 1.0 | 1.0 |
| **a1-2** | 1.0 | 1.0 |
| **a1-3** | 1.0 | 1.0 |
| **a1-4** | 1.0 | 1.0 |
| **a1-6** | 1.0 | 1.0 |
| **a2-8** | 1.0 | 1.0 |
| **a2-9** | 1.0 | 1.0 |
| **b2-3** | 1.0 | 1.0 |
| **b2-6** | 1.0 | 1.0 |
| **4dGal(b1-4)GlcNAc** | 1.0 | 1.0 |
| **4dNeu5Ac(a2-3)Gal** | 1.0 | 1.0 |
| **6dGal(b1-4)GlcNAc** | 1.0 | 1.0 |
| **7dNeu5Ac(a2-3)Gal** | 1.0 | 1.0 |
| **8dNeu5Ac(a2-3)Gal** | 1.0 | 1.0 |

| | | |
|---|---|---|
| **9dNeu5Ac(a2-3)Gal** | 1.0 | 1.0 |
| **Fuc(a1-2)Gal** | 1.0 | 1.0 |
| **Fuc(a1-2)GalOS** | 1.0 | 1.0 |
| **Fuc(a1-2)GlcNAc** | 1.0 | 1.0 |
| **Fuc(a1-3)Glc** | 1.0 | 1.0 |
| **Fuc(a1-3)GlcN** | 1.0 | 1.0 |
| **Fuc(a1-3)GlcOS** | 1.0 | 1.0 |
| **Fuc(a1-4)GlcNAc** | 1.0 | 1.0 |
| **Fuc(a1-6)GlcNAc** | 1.0 | 1.0 |
| **Fuc(b1-3)GlcNAc** | 1.0 | 1.0 |
| **Gal(a1-2)Gal** | 1.0 | 1.0 |
| **Gal(a1-3)Fuc** | 1.0 | 1.0 |
| **Gal(a1-3)Gal** | 1.0 | 1.0 |
| **Gal(a1-3)GalNAc** | 1.0 | 1.0 |
| **Gal(a1-4)Gal** | 1.0 | 1.0 |
| **Gal(a1-4)GlcNAc** | 1.0 | 1.0 |
| **Gal(a1-6)Gal** | 1.0 | 1.0 |
| **Gal(a1-6)Glc** | 1.0 | 1.0 |
| **Gal(b1-2)Gal** | 1.0 | 1.0 |
| **Gal(b1-3)Gal** | 1.0 | 1.0 |
| **Gal(b1-3)GlcN** | 1.0 | 1.0 |
| **Gal(b1-3)GlcNAc** | 1.0 | 1.0 |
| **Gal(b1-4)Fuc** | 1.0 | 1.0 |
| **Gal(b1-4)Gal** | 1.0 | 1.0 |
| **Gal(b1-4)GalNAc** | 1.0 | 1.0 |
| **Gal(b1-4)Glc** | 1.0 | 1.0 |
| **Gal(b1-4)GlcNAcOP** | 1.0 | 1.0 |
| **Gal(b1-4)GlcOS** | 1.0 | 1.0 |
| **Gal(b1-6)Gal** | 1.0 | 1.0 |
| **Gal(b1-6)GalNAc** | 1.0 | 1.0 |
| **GalNAc(a1-3)Fuc** | 1.0 | 1.0 |
| **GalNAc(a1-3)Gal** | 1.0 | 1.0 |
| **GalNAc(a1-3)GalNAc** | 1.0 | 1.0 |
| **GalNAc(a1-4)Gal** | 1.0 | 1.0 |
| **GalNAc(b1-3)Gal** | 1.0 | 1.0 |
| **GalNAc(b1-3)GalNAc** | 1.0 | 1.0 |
| **GalNAc(b1-3)GlcNAc** | 1.0 | 1.0 |
| **GalNAc(b1-4)Gal** | 1.0 | 1.0 |
| **GalNAc(b1-6)Gal** | 1.0 | 1.0 |
| **GalNAc(b1-6)GalNAc** | 1.0 | 1.0 |
| **GalNAcOS(b1-4)Gal** | 1.0 | 1.0 |

| | | |
|---|---|---|
| **GalNAcOS(b1-4)GlcNAc** | 1.0 | 1.0 |
| **GalNAcOS(b1-4)GlcNAcOS** | 1.0 | 1.0 |
| **GalOP(b1-4)GlcNAc** | 1.0 | 1.0 |
| **GalOS(b1-3)GalNAc** | 1.0 | 1.0 |
| **GalOS(b1-3)GlcNAc** | 1.0 | 1.0 |
| **GalOS(b1-3)GlcNAcOS** | 1.0 | 1.0 |
| **GalOS(b1-4)Glc** | 1.0 | 1.0 |
| **GalOS(b1-4)GlcN** | 1.0 | 1.0 |
| **GalOS(b1-4)GlcNAcOS** | 1.0 | 1.0 |
| **GalOS(b1-4)GlcOS** | 1.0 | 1.0 |
| **Glc(a1-4)Glc** | 1.0 | 1.0 |
| **Glc(a1-6)Glc** | 1.0 | 1.0 |
| **Glc(b1-4)Glc** | 1.0 | 1.0 |
| **Glc(b1-6)Glc** | 1.0 | 1.0 |
| **GlcA(b1-3)Gal** | 1.0 | 1.0 |
| **GlcA(b1-3)GlcNAc** | 1.0 | 1.0 |
| **GlcA(b1-6)Gal** | 1.0 | 1.0 |
| **GlcN(b1-3)Gal** | 1.0 | 1.0 |
| **GlcNAc(a1-3)Gal** | 1.0 | 1.0 |
| **GlcNAc(a1-4)Gal** | 1.0 | 1.0 |
| **GlcNAc(a1-6)Gal** | 1.0 | 1.0 |
| **GlcNAc(b1-2)Gal** | 1.0 | 1.0 |
| **GlcNAc(b1-2)GlcNAc** | 1.0 | 1.0 |
| **GlcNAc(b1-3)Fuc** | 1.0 | 1.0 |
| **GlcNAc(b1-3)GalNAc** | 1.0 | 1.0 |
| **GlcNAc(b1-3)GlcNAc** | 1.0 | 1.0 |
| **GlcNAc(b1-3)Man** | 1.0 | 1.0 |
| **GlcNAc(b1-4)** | 1.0 | 1.0 |
| **GlcNAc(b1-4)Gal** | 1.0 | 1.0 |
| **GlcNAc(b1-4)GalNAc** | 1.0 | 1.0 |
| **GlcNAc(b1-6)Gal** | 1.0 | 1.0 |
| **GlcNAc(b1-6)GlcNAc** | 1.0 | 1.0 |
| **GlcNAcOProp(b1-4)GlcNAc** | 1.0 | 1.0 |
| **GlcNAcOS(b1-3)Gal** | 1.0 | 1.0 |
| **GlcOSA(b1-3)Gal** | 1.0 | 1.0 |
| **Kdn(a2-3)Gal** | 1.0 | 1.0 |
| **Kdn(a2-6)Gal** | 1.0 | 1.0 |
| **Man(a1-2)Man** | 1.0 | 1.0 |
| **Man(a1-3)Gal** | 1.0 | 1.0 |
| **Man(a1-3)GlcNAc** | 1.0 | 1.0 |

| | | |
|---|---|---|
| **Man(a1-6)GlcNAc** | 1.0 | 1.0 |
| **MurNAc(b1-4)GlcNAc** | 1.0 | 1.0 |
| **Neu(a2-3)Gal** | 1.0 | 1.0 |
| **Neu(a2-3)GalOS** | 1.0 | 1.0 |
| **Neu(a2-6)Gal** | 1.0 | 1.0 |
| **Neu5Ac(a2-3)4dGal** | 1.0 | 1.0 |
| **Neu5Ac(a2-3)6dGal** | 1.0 | 1.0 |
| **Neu5Ac(a2-8)Neu5Ac** | 1.0 | 1.0 |
| **Neu5Ac(a2-8)Neu5Gc** | 1.0 | 1.0 |
| **Neu5Ac(a2-9)Neu5Ac** | 1.0 | 1.0 |
| **Neu5Ac(b1-6)GalNAc** | 1.0 | 1.0 |
| **Neu5Ac(b2-3)Gal** | 1.0 | 1.0 |
| **Neu5Ac(b2-6)Gal** | 1.0 | 1.0 |
| **Neu5Ac(b2-6)GalNAc** | 1.0 | 1.0 |
| **Neu5Ac9Ac(a2-3)Gal** | 1.0 | 1.0 |
| **Neu5Ac9Ac(a2-6)Gal** | 1.0 | 1.0 |
| **Neu5AcOMe(a2-3)Gal** | 1.0 | 1.0 |
| **Neu5Gc(a2-3)Gal** | 1.0 | 1.0 |
| **Neu5Gc(a2-6)Gal** | 1.0 | 1.0 |
| **Neu5Gc(a2-6)GalNAc** | 1.0 | 1.0 |
| **Neu5Gc(a2-8)Neu5Ac** | 1.0 | 1.0 |
| **Neu5Gc(a2-8)Neu5Gc** | 1.0 | 1.0 |
| **Neu5Gc(b2-6)Gal** | 1.0 | 1.0 |
